# Lateral hypothalamic melanin-concentrating hormone neuron dynamics in rats during sensory stimulation and sugar sweetened alcoholic cocktail drinking

**DOI:** 10.64898/2026.04.17.719280

**Authors:** Isabel R. K. Kuebler, Jonathan Y. Chin, Jessica D. Vollan, Mauricio Suarez, Caroline E. Bass, Nicholas A. Hubbard, Ken T. Wakabayashi

## Abstract

There is a dearth of information on how different cocktails sweetened with different sugars impact brain activity. Glucose enters the brain faster and in greater concentration than fructose and directly affects neuronal activity of melanin-concentrating hormone (MCH) neurons. MCH signaling promotes both glucose drinking and alcohol intake by integrating central and sensory inputs, but it is currently unknown how MCH neuronal activity relates to sweetened cocktail drinking. This study sought to investigate the relationship between MCH activity and sugar-sweetened alcoholic cocktail drinking. We also sought to compare MCH neuronal responses to the sugar solutions without alcohol as well as their response to sensory stimuli. In female and male rats, we used fiber photometry to monitor MCH neurons in response to sensory stimuli and during drinking of 10% glucose, 10% fructose, and glucose or fructose cocktails with 1.25% or 10% alcohol. We found that MCH activity rises in response to a variety of sensory stimuli and peaks before the start of drinking for all cocktails, before returning to baseline near the start of drinking. The cocktail type impacted the dynamics of MCH activity, where increased alcohol concentration resulted in earlier MCH activity for fructose but not glucose cocktails. Finally, we found that peak MCH activity during drinking is correlated with approach behavior for all sugar and cocktail types. These findings suggest that glucose and alcohol may interact to directly influence MCH activity. Further, MCH neurons may regulate cocktail drinking in response to sugar type and alcohol concentration.

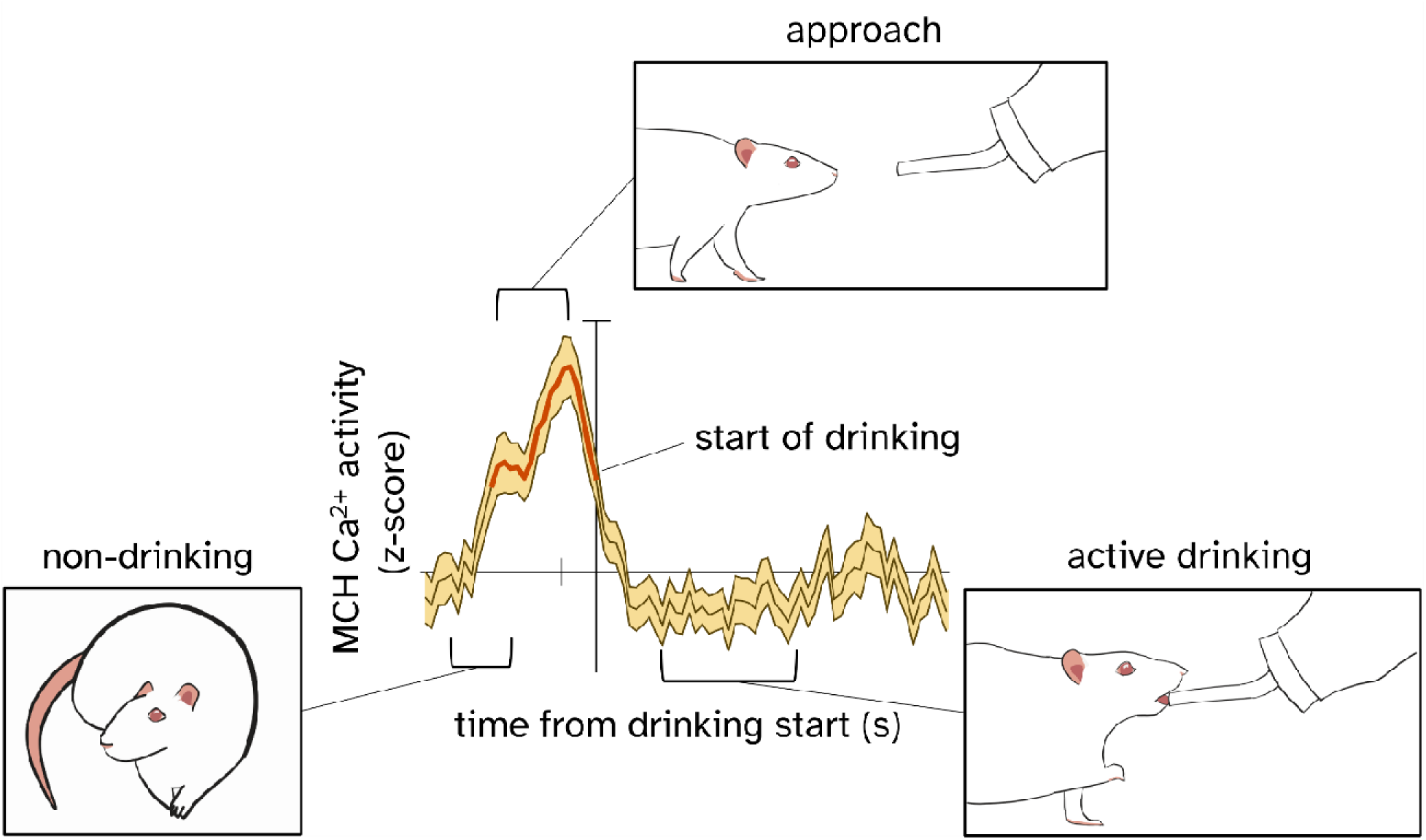

**New and noteworthy:** Fiber photometry was used to monitor lateral hypothalamic melanin-concentrating hormone (MCH) neurons in male and female rats during sensory stimuli and drinking of glucose, fructose, or glucose- or fructose-sweetened alcoholic cocktails. Subsecond-scale changes in MCH activity occurred after stimuli. Peak MCH activity during drinking was correlated with approach behavior. Alcohol concentration only impacted MCH activity with fructose cocktails. We discuss the implications of MCH dynamics towards brain function, associative learning, and alcohol use disorder.

## Introduction

Alcohol Use Disorder (AUD) is a major health problem. In the United States, 29.5 million Americans aged twelve and older self-report having symptoms of AUD (SAMHSA, 2023). Due to its high prevalence in the US, $249 million is spent each year on AUD including productivity loss, healthcare, and legal expenses (Sacks et al., 2015). Flavored alcoholic beverages (FABs), including “ready-to-drink cocktails”, are becoming more popular (NIQ, 2022) especially among youths (Albers et al., 2015; Fortunato et al., 2014; Griffiths & Sutherland, 1998). Many people start drinking during early adolescence (CDC, 2024), and frequent episodic drinking during adolescence corresponds to greater risk for AUD in adulthood (Patrick et al., 2023; Sloan et al., 2011). Therefore, understanding how sweetened cocktails could promote alcohol drinking may prove critical for developing effective prevention or treatment strategies for AUD.

Research on sweetened cocktail drinking has mostly focused on taste as the primary factor regulating drinking (Albers et al., 2015; Sardarian et al., 2020). Some studies have investigated additional factors including alcohol concentration (Jones & Reis, 2012) and caloric value (Ayoub et al., 2020) for cocktail drinking. However, research has not yet determined how different cocktail types impact the brain. Specifically, FABs often contain mixes of sugars such as high-fructose corn syrup, which is comprised of two monosaccharides, glucose and fructose (Khorshidian et al., 2021). Glucose is the brain’s main energy source (Duelli & Kuschinsky, 2001) and directly regulates the activity of some neurons (Ashford et al., 1990; Oomura et al., 1969). In contrast, while fructose can also enter the brain, fructose crosses the blood-brain barrier much slower and in much less concentration than glucose (Duelli & Kuschinsky, 2001). Additionally, there is currently no known mechanism by which fructose affects the activity of neurons in the mammalian brain. One neuronal system that changes its activity in response to brain glucose is the melanin-concentrating hormone (MCH) system.

In slice preparations, elevated extracellular glucose concentrations hyperpolarize MCH neurons, sensitizing them for increased neuronal firing (Burdakov et al., 2005). Rapid and large increases in blood glucose from oral intake or intravenous administration of glucose solutions are also tightly correlated with large increases in extracellular glucose concentrations in multiple brain areas including the lateral hypothalamus, where a large population of MCH expressing neurons are found (Kuebler et al., 2022; Wakabayashi et al., 2016; Wakabayashi & Kiyatkin, 2015). Thus, drinking glucose likely increases MCH neuronal activity in an intact brain, which may subsequently influence behavior.

The MCH system is thought to regulate energy balance by promoting feeding (Routh et al., 2014) as well as influencing a variety of motivated behaviors (Diniz & Bittencourt, 2017). Indeed, MCH receptor binding promotes both sucrose and glucose drinking (Benoit et al., 2005; Duncan et al., 2005; Sakamaki et al., 2005). Moreover, MCH signaling is necessary and sufficient for the preference of caloric sweeteners (i.e., sucrose) over non-caloric sweeteners (i.e., sucralose) (Domingos et al., 2013). Thus, the MCH system may respond to changes in central glucose levels by regulating feeding behavior, increasing the available energy sources for the brain.

One way that the MCH system may promote energy balance is by modulating sensory inputs to regulate behavior. For example, the MCH system rapidly integrates sensory information to regulate feeding and parental behaviors (for review see Diniz & Bittencourt, 2017; Kuebler et al., 2024). In support of this idea, we have previously shown when rats are briefly presented with a novel object or given a mildly irritating tail touch, extracellular glucose concentrations rapidly rise in the lateral hypothalamus in response to naturalistic stimuli of different modalities (Kuebler et al., 2022). Given that these changes in extracellular glucose concentrations are tightly correlated with changes in neuronal activity (Fellows et al., 2006; Hoge et al., 2005; Kiyatkin & Lenoir, 2012), MCH neurons are poised to respond to both energic and multimodal sensory information at behaviorally-relevant timescales. Others have shown that MCH neuronal activity is involved in distinguishing known and unknown stimuli such as a novel object (Gonzalez et al., 2016) or a brief auditory stimulus (Miller et al., 1993). Given these results, a plausible hypothesis is that MCH integrates both central glucose levels and external sensory stimuli to coordinate feeding behavior.

There are also strong links between MCH and alcohol drinking behavior. First, MCH regulation of alcohol drinking has been widely studied (Kuebler et al., 2024; Morganstern et al., 2020). For alcohol without any added sweeteners, MCH signaling promotes alcohol intake, where binding of MCH to MCH receptors throughout the brain increases alcohol drinking (Duncan et al., 2005). Conversely, decreasing MCH receptor function by MCH receptor antagonism or receptor knockout decreases alcohol seeking and intake (Cippitelli et al., 2010; Karlsson, Aziz, et al., 2016). Second, alcohol intake also impacts the expression of MCH in neurons. For example, short-term alcohol consumption is correlated with long-term increased MCH expression (Pickering et al., 2007). Further, the length of alcohol consumption is critical for MCH expression. Acute alcohol intake results in increased MCH expression, while chronic alcohol intake results in decreased MCH expression (Morganstern, Chang, Chen, et al., 2010). Despite these findings, the mechanisms of how MCH and alcohol drinking interact are still unknown.

Indirect evidence indicates a role for MCH neurons in regulating both sugar and alcohol seeking and intake. To wit, we recently examined the impact of alcoholic cocktails with either glucose or fructose on alcohol free drinking. We found a nuanced effect of sugar type on alcohol concentration: rats drank more alcohol per body weight with glucose-sweetened cocktails at lower alcohol concentrations while drinking more fructose-sweetened cocktails at high alcohol concentrations (Kuebler et al. 2026). Moreover, we found behavioral evidence that the post-ingestive central effects differed across glucose- and fructose-sweetened cocktails.

As one of the neuronal populations sensitive to central glucose levels, MCH neuronal activity could be related with sweetened alcoholic cocktail drinking. Yet there is little direct evidence for how MCH activity is associated with ingestive behavior such as sweetened alcoholic cocktail drinking. Traditional methods such as systemic manipulations such as MCH receptor antagonism (Karlsson, Aziz, et al., 2016) lack the temporal precision necessary to elucidate how MCH neuronal activity is correlated with individual feeding or drinking behaviors. Further, some recording methods with greater temporal resolution such as electrophysiology cannot provide neurochemical specificity *in vivo* (van den Pol et al., 2004). Given the limitations of these typical methods, it is unknown how MCH neurons respond to alcohol drinking behavior when it is mixed with sugars of different central penetrance. This presents a considerable gap in our understanding because it may be that unique patterns of MCH neuronal activity are correlated with different phases of cocktail drinking. For example, it is possible that raised MCH neuronal activity in the lateral hypothalamus is associated primarily with the appetitive phase of drinking, including approach behaviors, or it may be associated more with the consummatory phase of active drinking (for review see Kongstorp et al., 2025).

Thus the central aim of this study was to determine relationship between MCH activity, their response to different sensory stimuli of different modalities, and alcohol cocktail drinking behavior. We used fiber photometry to elucidate these relationships, as this approach permits sufficient neuronal specificity and temporal resolution to resolve this question. This newer technique allowed us to selective monitor MCH neurons in the lateral hypothalamus with subsecond precision to distinguish between phases of drinking behavior. Additionally, fiber photometry permitted repeated recordings from the same animals over extended periods of time.

Our second aim was to determine whether MCH activity differed among different cocktails and between alcohol-free glucose and fructose drinking. Specifically, we investigated the timing and magnitude of MCH activity during drinking and sensory stimuli, and the impact of different sugar and cocktail types. We also sought to ascertain whether both female and male rats responded to each stimulus and to establish experimental procedures in both sexes. We hypothesized that increases in MCH neuronal activity are correlated with sensory stimuli of different modalities including novel object presentation, tail touch, a brief auditory stimulus, and ambient light changes. Germane to drinking behavior, we predicted that MCH neuronal activity would rise at the start of drinking and return to baseline during drinking. Further, we predicted that the rise in MCH activity would be potentiated by the entry of glucose and alcohol, but not fructose, into the brain. During the experiment, we discovered that the dynamics of MCH activity were more complex than predicted, and so a final objective of this study was to determine which phases of drinking were most associated with changes in MCH activity for each sugar or cocktail solution.

## Methods

### Animals

Data were collected from male (n=2) and female (n=3) adult Wistar rats, weighing 270-290g (male) and 250-270g (female) upon arrival. Average ages were 3.9 months (males) and 4.9 months (females) at the start of experimental recordings and 6.5 months (males) and 8.0 months (females) at the end of recordings. Animals were housed in pairs in HVAC filtered home cages upon arrival and singly starting the day before surgery, in a climate-controlled colony with standard chow (Teklad) and tap water *ad libitum* in their home cages. Rats were maintained on a reverse light-dark cycle, with lights on starting at 1500. Rats were acclimatized to the animal facility for at least 7 days before surgical procedures. All subjects were intact, and female subjects were free-cycling, as we had no *a priori* hypotheses on the effect of ovarian cycles on MCH activity. All animal procedures were approved by the University of Nebraska-Lincoln Institutional Animal Care and Use Committee and abide by the Guide for the Care and Use of Laboratory Animals (National Research Council (U.S.) et al., 2011).

### Combinatorial virus production

We have previously used two adeno-associated viruses (AAVs) in an intersectional approach to target transgene expression to neuronal subtypes including dopamine, γ-aminobutyric acid (GABA) neurons, and MCH neurons (Gompf et al., 2015; Suarez et al., 2026; Wakabayashi et al., 2019). In order to target MCH neurons, the LH was co-infused with an AAV that expresses Cre recombinase under the control of a minimal MCH promoter (MCH-Cre-AAV) and a Cre-dependent virus (DIO-gCaMP6f), so that only MCH neurons expressing Cre would flip the gCaMP6f transgene to the sense orientation, resulting in GCaMP6f expression only in MCH neurons.

A 461 bp 5’ element upstream from the MCH coding sequence was identified in the NCBI database (accession number M62641.1, Thompson & Watson, 1990). This minimal promoter sequence has previously been used to drive expression in MCH neurons in the LH (Blanco-Centurion et al., 2016; Suarez et al., 2026). We noted a one-base discrepancy (an additional A after base 354 of the reported 462 base promoter in (Blanco-Centurion et al., 2016)) which we identified as a typographical error based on alignment to the M62641.1 sequence. A transgene consisting of the MCH promoter, 2nls-iCre (“improved Cre” with a nuclear localization signal), P2A sequence, mCherry, and a beta-globin intron was synthesized (GeneArt, Thermo Fisher Scientific, Waltham, MA). This transgene was digested at the 5’ *Nhe*I and 3’ *Xho*I sites of the gene synthesis product, and cloned into corresponding 5’ *Nhe*I and 3’ *Sal*I sites of the pAAV-prMCS-iCre plasmid (original construct generated in the Bass laboratory), which effectively replaces the original β-globin intron-iCre transgene. The resulting plasmid was named MCH-2nls-iCre-P2A-mCherry-pAAV and packaged in the AAV10 serotype of AAV using the standard triple transfection protocol in HEK-293 cells (Xiao et al., 1998) to create recombinant pseudotyped AAV2/10 virus (titer:1.491014 vector genome copies per mL). Hereafter we will refer this viral construct as MCH-Cre-P2A-mCherry for clarity. We also packaged Ef1a-DIO-gCaMP6f-TdTomato-AAV2/10 (Addgene #51083; titer: ≥1x10^12^ vcg/mL), hereafter called gCaMP6f. These were co-infused, 125 nL of MCH-Cre-P2A-mCherry + 375 nL gCaMP6f, 500 nL total per side, into the LH, resulting in gCaMP6f expression restricted to MCH neurons.

### Stereotaxic surgery

Rats were given pre-operative carprofen (Bio-serv, 5mg/kg, oral tablet, MD150-2) no more than 24 hours before surgery. General anesthesia was induced with a ketamine HCl (33 mg/kg) and xylazine (10 mg/kg) mix intraperitoneally, and anesthesia was maintained with inhaled isoflurane (1-2%, SomnoSuite, Kent Scientific, VetOne) using either room air and an oxygen condenser (DeVilbiss, Port Washington, NY), or an oxygen tank. 500 nL of virus was injected bilaterally using a microinjector (Drummond Scientific, Broomall, PA) to the lateral hypothalamus: AP -3.0 mm, ML +/- 1.6 mm, DV -8.8mm according to the stereotaxic atlas of Paxinos and Watson (2014), similarly to our recent work (Suarez et al., 2026). Post-infusion wait time until removing the injector was 5 minutes. Fiber optic cannulae (MBF Biosciences, Williston, VT, FOC-BF-200-125) were implanted bilaterally to the dorsal area of the lateral hypothalamus: AP -3.0 mm, ML +/- 1.6 mm; DV -7.8 mm (**Fig. 1a**). The skull was prepared with C&B-Metabond etchant and primer (Parkell, Edgewood, NY) before viral infusions, and fiber optic cannulae were secured with dental acrylic (Lang Dental Mfg. Co., Wheeling, IL) anchored by three stainless steel bone screws (BASi, West Lafayette, IN, MD-1310). Post-operative analgesia 5 mg/kg carprofen was administered s.c. (Rimadyl, Zoetis, Parsippany, NJ) immediately after the end of procedures, and daily carprofen 2mg oral was given two days post-surgery. Experimental procedures started ∼28 days after virus infusion to allow for virus expression.

**Fig. 1.**
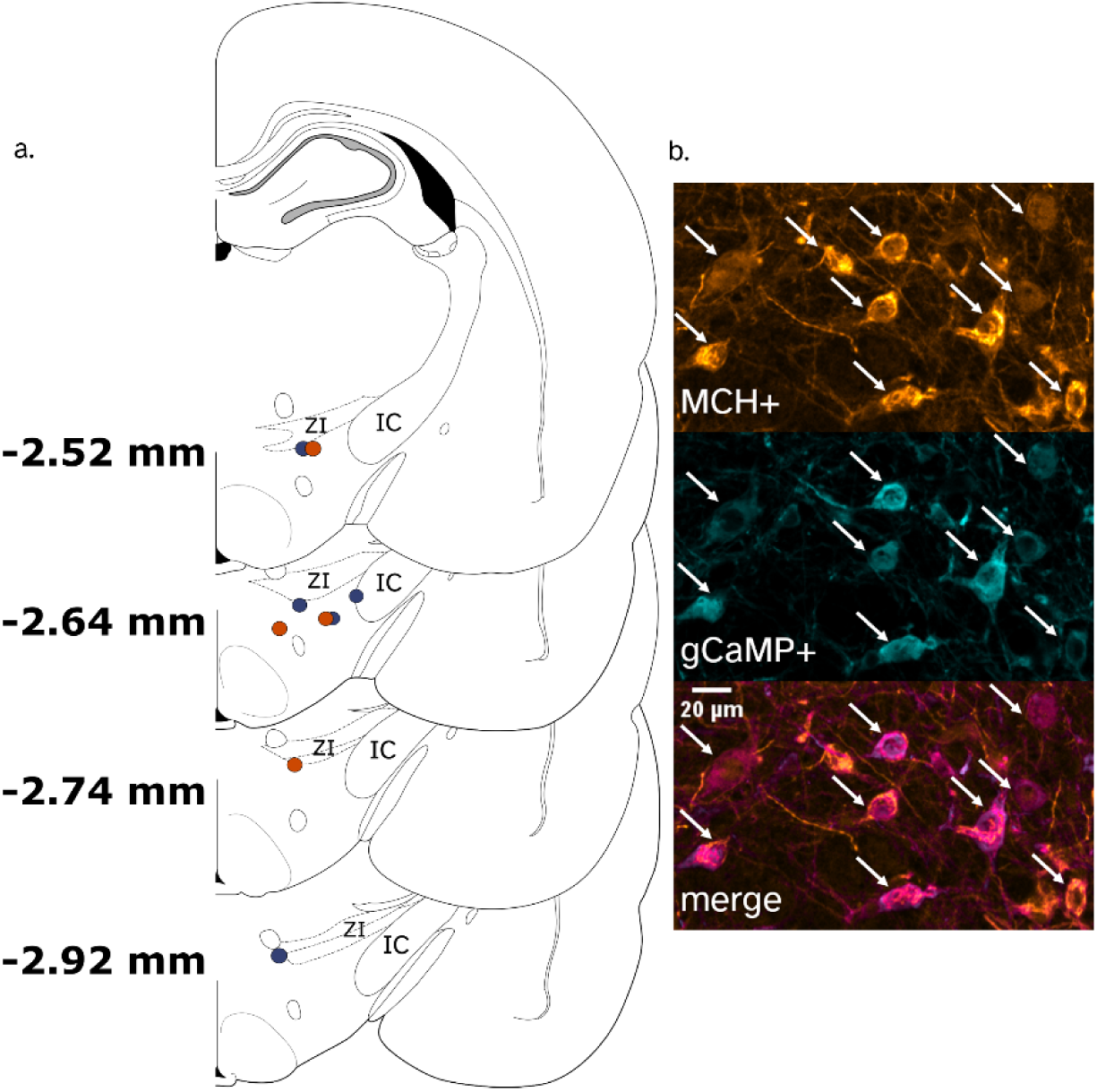
Histological verification of cannula placement and viral specificity. (**a**) Schematic of tip placement of the fiber optic cannulae in the lateral hypothalamus. Note that the recording area is ventral to the cannula tip. Female subject placements are denoted in blue, and male subject placements are denoted in orange. All fiber optic cannulae used in analyses were ventral to the zona incerta (ZI), demonstrating selective recording from the lateral hypothalamus. Atlas figures adapted from (Paxinos & Watson, 2006) (**b**) Representative image of MCH gCaMP6f expression. MCH-positive neurons (orange) showed selective expression of gCaMP6f-positive neurons (cyan) shown by colocalized neurons (magenta).

### Calcium imaging

We performed fiber photometry using the FP3002 system running Bonsai workflows (MBF Biosciences, Williston, VT). A 2-branching patch cord, 2.5 meters long (Doric Lenses, Québec City, Canada, BFP(2) 200/220/900-0.37 2.5m_FCM*-2xMF1.25) was used. We used one wavelength of LED light to excite the calcium-dependent gCaMP6f fluorescence (470 nm), another that excited isosbestic fluorescence from other sources (415 nm) to control for autofluorescence and movement artefacts, and a third to capture native fluorescence from the tdTomato tag on the gCaMP6f (560 nm, not used in this analysis). LED light sources alternated continuously (no off time) among 415 nm (isosbestic), 470 nm (Ca^2+^-dependent), and 560 nm (td-Tomato) wavelengths. Patch cords were photobleached for 12 hours before the start of each cohort to inhibit fluorescence within the patch cord itself. Alignment of the patch cords to the FP3002 camera and regions of interest (ROIs) were confirmed before recording each day. Bonsai workflows were set up in the same way for each recording: raw fiber photometry data were captured and saved as a .csv file, and the timestamps of experimenter keypresses were recorded.

### Recording protocol

A custom 24 cm w x 30 cm l x 30 cm h clear acrylic chamber (Central Plasticworks, Lincoln, NE) with bottle nozzle holes 8 cm from the base and 4 cm from the sides was used for recording. The chamber sat atop a 28 cm diameter turntable (Wilton, Naperville, IL) for manual turning to reduce coiling of the patch cord. Fresh bedding (Teklad) was used for each subject, and the chamber was cleaned with Caviwipes (Metrex, Romulus, MI) and air dried between each subject. Rats were acclimatized to the recording chamber once for ∼ 1 hr at least 1 day before recording sessions.

On the day of a recording, rats were gently restrained using a hand towel, connected to the patch cord with a ceramic sleeve (MBF Biosciences, Williston, VT, SM-CS125S), placed in the recording chamber, and left for 15 minutes to acclimate to the chamber before experimental stimuli. The rat was recorded during this time to verify the connection between the patch cord and the optical ferrule as well as the overall reliability and stability of the recording (data not shown). Recording sessions were capped at 15 min, after which the LEDs were turned off for 5 minutes to minimize photobleaching.

### Solutions

Sugar and sugar-sweetened alcohol solutions were mixed using filtered tap water and either D-(-)-fructose (Sigma-Aldrich, St. Louis, MO) or D-(+)-glucose (Oakwood Chemical, Estill, SC). Each solution had 10% glucose or fructose (w/v) alone, or with 1.25% or 10% ethanol added (v/v), using 95% ethanol (Decon Labs, King of Prussia, PA). Plastic drinking bottles containing a ball valve (MedAssociates) were weighed before and after the session to measure the volume drank.

### Sensory stimuli

Each rat was exposed to several sensory stimuli to assess the MCH neuronal response. Tail touch was conducted last (following all other sensory stimuli) to minimize the possibility of carryover stress. All non-drinking stimuli recordings were conducted before drinking trials to minimize putative contributions of central post-ingestive effects on neuronal responses to the stimuli.

### Novel Object

After 5 minutes of baseline recording, a novel object was placed in the chamber by the experimenter similarly to (Kuebler et al., 2022): either a 25 ml glass vial with a white plastic cap, a multicolored plastic chemical model of alcohol, or a black plastic fork. Each subject received the glass vial for three recordings, and any subsequent recordings included a different novel object. The novel object was removed after 1 minute, and the timestamps of the experimenter placing the object on the floor of the chamber and picking up the object were recorded using keypresses. Recording stopped 5 minutes after the end of the stimulus, and the LED was turned off for 5 minutes.

### Tail touch

After 5 minutes of baseline, an experimenter manually placed a wooden clothespin on the rat’s tail approximately one-third to one-half the length of the tail from the distal end. As the clothespin had a hole around the size of the rat’s tail, this provided a mildly arousing stimulus. 1 minute later the clothespin was removed. Timestamps of either the clothespin being placed on the tail or the experimenter taking hold of the tail, and the removal of the clothespin, were recorded using keypresses. Recording stopped 5 minutes after the end of the stimulus, and the LED was turned off for 5 minutes.

### Room light

After 2.5 minutes of baseline, the room light was turned off, then turned back on 10 seconds later. Timestamps of light switches were recorded using keypresses. Recording stopped 2.5 minutes after the end of the stimulus, and the LED was turned off for 5 minutes.

### Auditory stimulus

After 2.5 minutes of baseline, a brief (0.5 s) auditory stimulus of 75 dB was given, and the auditory stimulus timestamp was recording using a keypress. Recording stopped 2.5 minutes after the end of the stimulus, and the LED was turned off for 5 minutes.

### Sugar and sweetened alcohol drinking

Before recording sessions, rats were habituated to drinking sugar solutions until they displayed robust drinking behavior: all rats were habituated to drinking 10% glucose once, and two female rats were also habituated to drinking 10% fructose once. All habituation sessions were conducted at least 1 day before recording sessions.

Across the recording sessions, the two solutions without alcohol (10% glucose and 10% fructose) were presented first, and the cocktail solutions were presented in order of increasing alcohol concentration, with the order of fructose and glucose cocktails randomized. Rats had no prior experience with 1.25% alcohol cocktails or with 10% glucose + 10% alcohol cocktails. With 10% fructose + 10% alcohol cocktails, 40% of recordings used in analyses were after previous experience drinking the same cocktail, as drinking behavior was less robust for this cocktail.

On the day of a recording, after 1 minute of baseline, the bottle nozzle was inserted into the chamber via the hole in the wall, and this event was timestamped using a keypress. Within the drinking session, the start and end of each drinking bout were timestamped by an experimenter. Data for male rats were analyzed using only keypress data. During the course of the experiment, we began video recording the sessions, first using an overhead camera (Stopmotion Explosion PRO HD 1080P) and later a close-up camera (GoPro Hero 5) affixed to the side of the chamber, facing the nozzle. Data from female rats were analyzed using video recordings. Video timestamps were used to validate the precision of data from keypresses. The bottle was removed 24 minutes after the start of drinking; this event was timestamped using a unique keypress, and the recording was stopped seconds after the bottle was removed.

### Histological verification

Subjects were deeply anesthetized with 390 mg/kg pentobarbital sodium (FatalPlus, Vortech Pharmaceuticals, Dearborn, MI) i.p. then transcardially perfused with a 400 mL ice cold perfusion rinse containing 100 mM phosphate buffered saline (PBS, Electron Microscopy Sciences, Hatfield, PA) containing 18 mM procaine HCl, and 73 mM sucrose followed by 250 mL of ice cold 10L% formalin (Sigma Aldrich), with a final pH of 7.4 (Suarez et al., 2026; Wakabayashi et al., 2010). Brains were stored in 10% formalin for 24 hours and then transferred to a solution of 30% sucrose and 100 mM PBS for at least 3 days. Brain tissue was sectioned at 35 μm with dry ice on a sliding microtome (American Optical, Buffalo, NY, model 860) and stored in cryoprotectant solution (Watson et al., 1986) at -20°C. To validate the placement of the fiber optic cannulae, floating sections were imaged on a custom inverted fluorescence microscope (Olympus) with a 4x LCPlanFl objective, using an INFINITY8-C model camera with Infinity Analyze software (version 7, Teledyne Solutions). Images were captured in brightfield, and two experimenters validated placement of the fiber optic cannulae (**Fig. 1a**).

Viral expression was further verified using subjects from this study (n=2) and subjects that were not implanted with fiber optic cannulae (n=2) to confirm that the double-floxed inverted open reading frame (DIO) transgenes were expressed in MCH+ neurons only. Expression of the gCaMP6f in MCH+ neurons was determined using immunohistochemistry as previously described (Suarez et al., 2026). First, sections were washed in 0.1 M Tris-buffered saline (TBS 1X, Growcells.com) for 30 minutes, pre-blocked with Image-iT FX Signal Enhancer in TBS + 0.5% Triton X-100 for 30 minutes, then pre-blocked in 5% normal donkey serum and 1% bovine serum albumin in TBS + 0.5% Triton X-100 for 2 hours at 23 °C. Next, sections were incubated with freshly prepared primary antibodies: rabbit anti-Melanin-Concentrating Hormone (MCH, 1:1000, H-070-47, Phoenix Pharmaceuticals) and goat anti-Green Fluorescent Protein (GFP, 1:1000, NB100-1770, Novus) at 23 °C for 6 hours and overnight at 4 °C. After washing in TBS, sections were incubated with freshly prepared compatible secondary antibodies: Donkey Cy5 anti-rabbit (1:400, 711-175-152, Jackson) and Donkey anti-goat AlexaFluor 488 (1:2000, A11055, Invitrogen) for 2 hours at 23 °C. Sections were mounted and coverslipped using Fluoromount-G (SouthernBiotech, Birmingham, AL) and high-magnification, high-resolution images were capture using a Nikon A1R inverted confocal microscope. The MCH/Cy5 signal was imaged at 640 nm, and the gCaMP6f/AlexaFluor signal was imaged at 488 nm. Z-stacks were imaged at 1.5 μm thickness at 20X magnification.

Images were postprocessed and analyzed using FIJI (Fiji is Just ImageJ), and quantification of MCH with gCaMP6f was performed by two experimenters (IRKK and KTW) similar to previously reported (Suarez et al., 2026). First, section images were chosen based on the mean intensity profile, so that slices used for quantification had maximum fluorescent intensity for both MCH and gCaMP6f antibodies. Next, images were manually adjusted for brightness and contrast, then assigned lookup tables for MCH (orange hot) and gCaMP6f (cyan), and the Colour merge function (MBF, n.d.) was applied for visual quantification of colocalized neurons (magenta). Neurons expressing MCH, gCaMP6f, or both were counted using the Cell Counter plugin (1008 total neurons, n=4). 63.94% of MCH+ neurons displayed colocalization of gCaMP6f, and 0.40% of neurons displayed gCaMP6f with unclear colocalization with MCH expression (**Fig. 1b**), which is similar to what others have reported when using the MCH promoter to drive transgene expression (Blanco-Centurion et al., 2016; Suarez et al., 2026).

### Data analysis

Data were analyzed using R version 4.3.3, FiPhA version 0.0.1 (Bridge et al., 2024), GraphPad Prism version 10.6, and JASP version 0.95.4.0.

### Preprocessing

Raw data were deinterleaved using an R script to separate the isosbestic, calcium-dependent, and td-Tomato wavelengths.

Preprocessing followed the procedure detailed in Martianova et al., 2019, with the exception of the current correction for photobleaching and normalization. Data for the isosbestic and calcium-dependent traces were analyzed in the same way using FiPhA: raw traces were subsetted to remove the first 100 rows of data (∼18s), detrended using a monoexponential curve, and standardized using z-scores to scale the isosbestic and calcium-dependent data to each other (**Fig. 2**). The isosbestic trace was further scaled to the calcium-dependent curve using linear regression in R, then the isosbestic signal was subtracted from the calcium-dependent signal to recover the signal without movement artefacts. Data from different subjects were analyzed without normalizing to avoid obscuring possible effects of sugar type or alcohol concentration (Wallace et al., 2025). A drinking bout was considered separate from the previous bout if it was separated by >= 1s, as finer precision was not consistent for real-time keypresses.

**Fig. 2.**
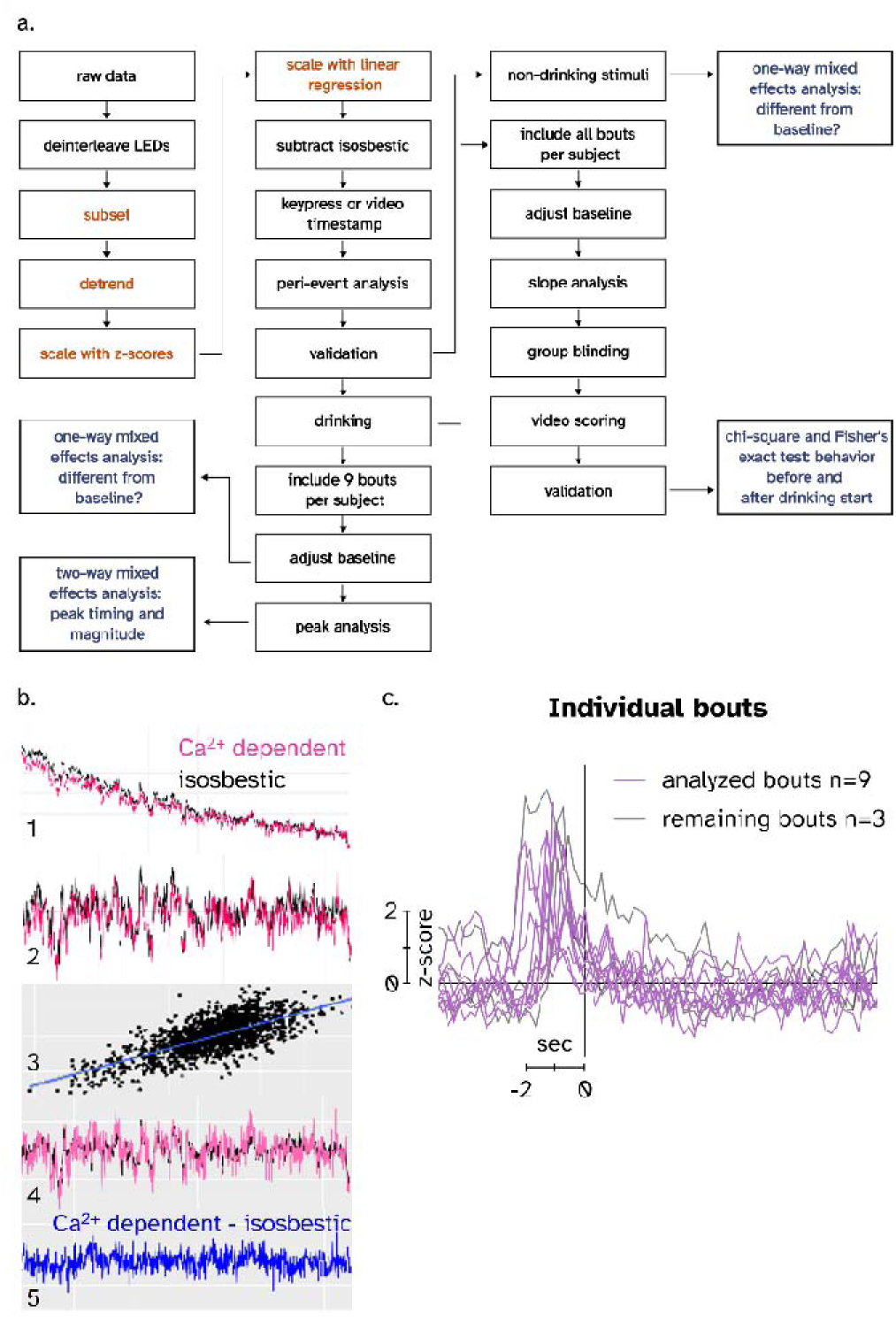
Data analysis workflow. (**a**) Schematic of data analysis. Orange text denotes analysis steps performed on both Ca^2+^ dependent and isosbestic traces separately. Blue text denotes statistical analyses. (**b**) Example preprocessing. 1. Subsetted Ca^2+^ dependent and isosbestic traces. 2. Removal of photobleaching. 3. Linear regression of standardized traces. 4. Linearly scaled Ca^2+^ dependent and isosbestic traces. 5. Isosbestic-subtracted trace.

### Validation

One rat had off-target fiber placement unilaterally, and another rat had no detectable signal for drinking behavior unilaterally; these data were removed from further analyses. All recordings were carefully examined by two experimenters for recording stability. Any sessions identified as having unreliable signal were carefully inspected further by three experimenters. We reviewed the experimental notes taken during the fiber photometric session, reassessed the recording for any possible unaccounted-for artefacts, and confirmed the stability of recordings for inclusion. The data was considered validated once the three raters came to a consensus on inclusion for each session. Sensory stimuli analyses included multiple recordings from each subject, while drinking analyses included only one recording per subject.

### Peri-event analysis

Thirty seconds before and after the start of each sensory or drinking event were included for peri-event analyses using FiPhA, and data were not normalized. Initial data preparation was completed using all bouts within the session for each subject and side of the brain contributing separately, all averaged together. As drinking behavior was naturalistic and not constrained, individual rats had different numbers of bouts per recording session. To ensure that all rats equally contributed to analysis, each rat contributed nine drinking bouts for each side of the brain. These bouts included the three bouts after the first bout to avoid artefacts during bottle insertion, the middle-most three bouts numerically, and the last three bouts of the session.

There were variable intervals between the earliest bouts and later bouts used in those analyses (2.3-22.9 min) and large differences in the amount of solution drank among sessions, so we chose not to compare early versus late drinking bouts.

Data for each recording were baseline-adjusted depending on the type of stimulus without normalizing. The start of drinking, novel object, room light, and auditory stimulus were adjusted by subtracting the average of -5 to -3s before the stimulus (0s). Tail touch was adjusted by subtracting the average of -7 to -5s before the stimulus, as the start of the procedure took several seconds longer than other stimuli.

The peak height, time of peak, and area under the curve (AUC) were analyzed in the same way for each drinking bout (**Supplemental Fig. S1a**). Peaks and respective AUCs were calculated for the entire +/- 30s peri-event window, and the peak closest to 1s before the start of drinking was used for the peak height analysis. AUC for this positive peak was recorded for the same peaks used for the peak height analysis. The y-value at drinking start was also found for each bout.

For each measurement of drinking stimuli, data from the nine bouts from each subject and side of the brain were combined for two-way Mixed Effects Analyses of sugar type and alcohol concentration. For non-drinking stimuli, all sessions from each subject and side of the brain were combined. As the groups were not powered to analyze for sex differences, we combined female and male data for all analyses. For each cocktail or non-drinking stimulus, the time points when the MCH signal was different from baseline (0 z-score) were analyzed as a single group using a one-way Mixed Effects Analysis of time, followed by a Fisher *post-hoc* test comparing each time point to baseline.

### Behavioral analysis

During the course of the experiment, we detected that the MCH signal rose before the start of drinking, peaked, and fell slowly until rising again. We chose to further analyze the average of all bouts in each drinking session (i.e., for each subject and each cocktail separately) as a descriptive ethology. First, the timestamp of the rise before the start of drinking was determined for each drinking session by finding the first derivative of the average of all drinking bouts and noting the earliest consistently positive value before the peak (**Supplemental Fig. S1b**). Then, the timestamp of the peak was found as part of the peak analysis described above. Lastly, the nadir when MCH signal starts to rise again was found using the first derivative, as the earliest instance of multiple consecutive positive values after the peak. If the two sides of the brain differed in the timestamps of these MCH signal changes, the timestamps from the side of the brain with more definite changes in slope were used to analyze data from both sides of the brain.

As only female subjects were consistently video recorded, ethology analysis was conducted on only females. Each drinking session resulted in varying numbers of bouts, so to equally represent each subject and cocktail, we again analyzed the same nine bouts per session: three bouts from the start of drinking (starting with the second bout to avoid the stimulus of the bottle being introduced into the chamber), three bouts from the middle of all the bouts, and the last three bouts. Two raters analyzed each video at the timestamps calculated for each change in slope of the MCH signal and recorded the behavior at that moment. Raters were blinded to the cocktail type but were not blinded to the timepoint, to ensure accuracy of the timestamps analyzed. The behaviors observed were selected from a list of behaviors shown in **Supplemental Table S1**, organized into several categories of “approach”/appetitive, “active drinking”/consummatory, drinking-related, and all other behaviors. First, a variety of appetitive behaviors were grouped together, including locomotion toward the nozzle (approach), movement of the head toward the nozzle while near the nozzle, and sniffing the nozzle (Mahler & Berridge, 2009; Tong et al., 2011). The category of drinking-related behaviors includes behaviors associated with hedonic or aversive responses, as well as interacting with the nozzle in ways other than sniffing or licking. Using overhead camera recording, it was not always possible to distinguish certain behaviors within a behavioral category. Here, rhythmic mouth movements and rhythmic tongue protrusions were categorized together, as they often occur in sequence (Grill & Norgren, 1978). The behavior of biting the nozzle refers to the rat taking hold of the nozzle without licking, which appeared to be an attempt to adjust the positioning of the nozzle or ball and was not clearly appetitive or consummatory. All behaviors outside of appetitive, consummatory, and drinking-related categories were grouped together. We did not distinguish general non-approach movement from movement away from the nozzle; therefore, this analysis does not specifically categorize avoidance behaviors resulting from aversive tastes (Schier & Spector, 2019).

The percentage of timestamps for which the two raters selected the same behavioral category was 86.0%. Any timestamps that were not in agreement between the two raters were reviewed by a group of three experimenters, and the group reached a final consensus on the behavior. Categorical variables were applied for each behavioral category using a custom R script, and the frequency of each behavior category was found for each cocktail type and each of the three timepoints around the start of drinking. A chi-square analysis was performed for each cocktail type of the frequencies of each behavior category across the three different timepoints. Exploratory *post-hoc* comparisons were performed between 1.25% and 10% alcohol for each sugar type, and between 10% glucose + 10% alcohol and 10% fructose + 10% alcohol using Fisher’s exact test. Active drinking versus all other behaviors were compared across cocktail type for the initial rise in MCH activity and for the nadir of MCH activity, while appetitive behaviors versus all other behaviors were compared across cocktail type for the peak of MCH activity.

## Results

### MCH calcium activity rises in response to a variety of sensory stimuli

To investigate when MCH neuronal activity changes in response to sensory stimuli, we first analyzed variability of MCH calcium activity across time for each of the sensory stimuli before investigating individual timepoints in *post-hoc* analyses. First, MCH calcium activity varied throughout the introduction of a novel object (**Fig. 3a**, one-way Mixed Effects Analysis, *F*_70,1750_=12.66, *p*<0.0001). MCH activity also varied during a tail touch (**Fig. 3b**, *F*_80,2160_=9.018, *p*<0.0001). Similarly, MCH activity varied across time when the room light was turned off (**Fig. 3c**, *F*_69,1656_=5.944, *p*<0.0001). Lastly, MCH activity varied throughout time with an auditory stimulus (**Fig. 3d**, *F*_70,2170_=16.57, *p*<0.0001). For both the novel object and tail touch stimuli, the MCH signal was significantly greater than baseline 1-2s before the start of the stimulus and more the majority of the 10s after the start of the stimulus. For the room light stimulus, the MCH signal became significantly greater than baseline immediately upon the start of the stimulus and remained elevated for much of the duration of the stimulus. Lastly, for the auditory stimulus, the MCH signal was significantly greater than baseline immediately in response to the stimulus, higher than for all other stimuli, and returned to baseline several seconds later.

**Fig. 3.**
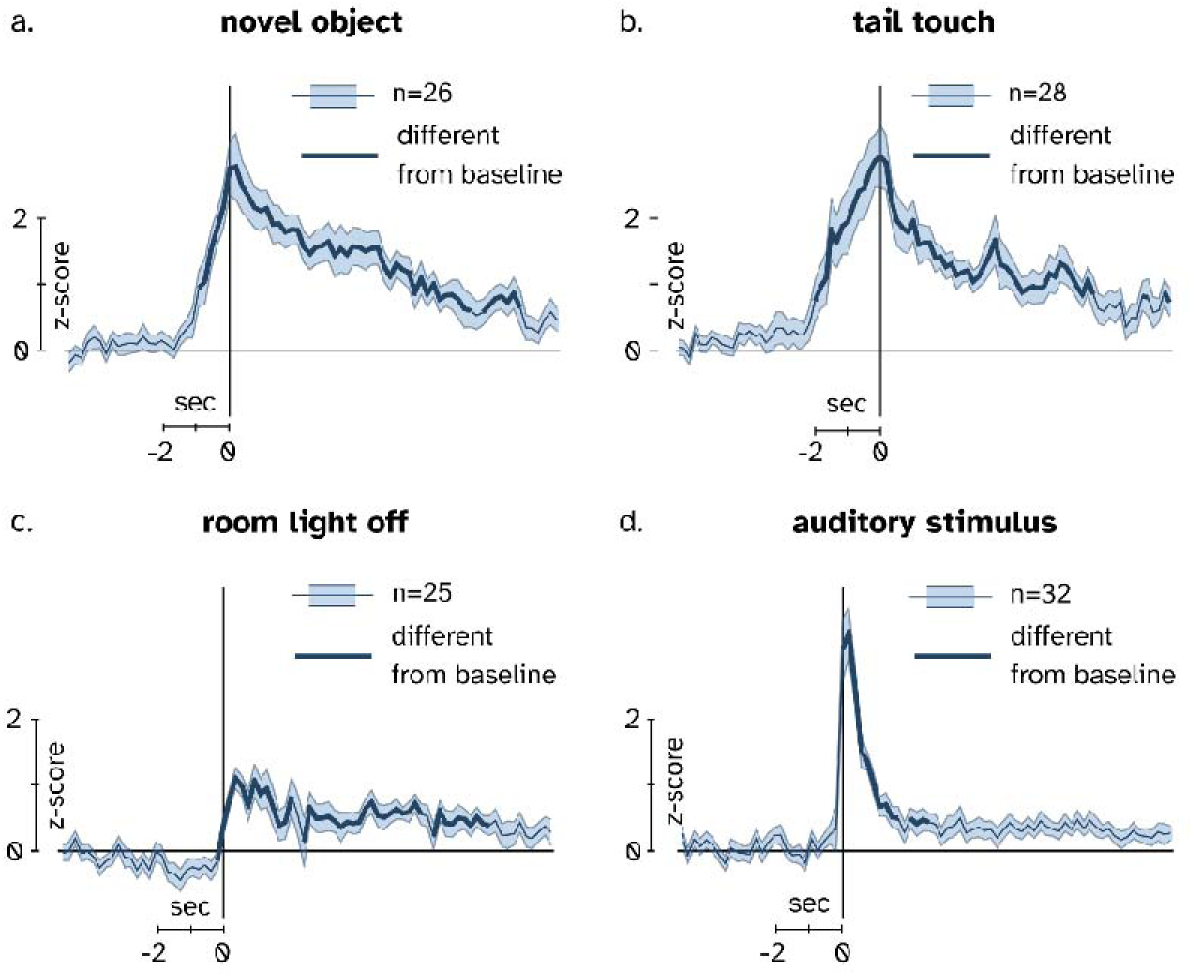
MCH activity rises at the start of each non-drinking stimulus. All data are shown as mean ± SEM. Y-axis scale bar denotes the 2 z-scores of MCH Ca^2+^ activity from baseline, X-axis scale bar denotes 2 seconds. Time intervals where MCH activity is different from baseline (0 z-score) are shown as a darker line on the mean (p<0.05). (**a**) MCH activity rose while the novel object was being placed, peaked when the novel object had been placed in the chamber (x=0), and slowly decreased back to baseline over 9 seconds. (**b**) MCH activity rose while the clothespin was being applied, peaked when the clothespin was placed on the tail, and slowly decreased toward baseline throughout the 10s after the clothespin was applied. (**c**) MCH activity quickly rose a modest amount when the room light was turned off and slowly decreased back to baseline over 8 seconds. (**d**) MCH activity quickly rose to the greatest magnitude of any stimulus in response to a brief auditory stimulus, quickly decreased within 1 second, and returned to baseline after 3 seconds.

### MCH calcium activity rises before the start of drinking and decreases after the start of drinking

To analyze when MCH activity changed during drinking behavior, we first analyzed whether MCH activity varied within the analysis window. MCH activity significantly varied from baseline throughout 10% glucose drinking (**Fig. 4a**, *F*_70,4970_=11.84, *p*<0.0001), 10% glucose + 1.25% alcohol drinking (**Fig. 4b**, *F*_70,4970_=7.919, *p*<0.0001), 10% glucose + 10% alcohol drinking (**Fig. 4c**, *F*_70,4970_=9.744, *p*<0.0001), 10% fructose drinking (**Fig. 4d**, *F*_70,4970_=11.11, *p*<0.0001), 10% fructose + 1.25% alcohol drinking (**Fig. 4e**, *F*_70,4970_=8.986, *p*<0.0001), and 10% fructose + 10% alcohol drinking (**Fig. 4f**, *F*_70,4970_=4.346, *p*<0.0001). For each sugar and cocktail, MCH calcium activity was significantly above baseline (-5 to -3s before the start of drinking (0s)) for several seconds before the start of drinking (**Fig. 4a-e**). Additionally, for cocktails containing 1.25% alcohol, MCH calcium activity was significantly below baseline after the start of drinking (**Fig. 4b, e**).

**Fig. 4.**
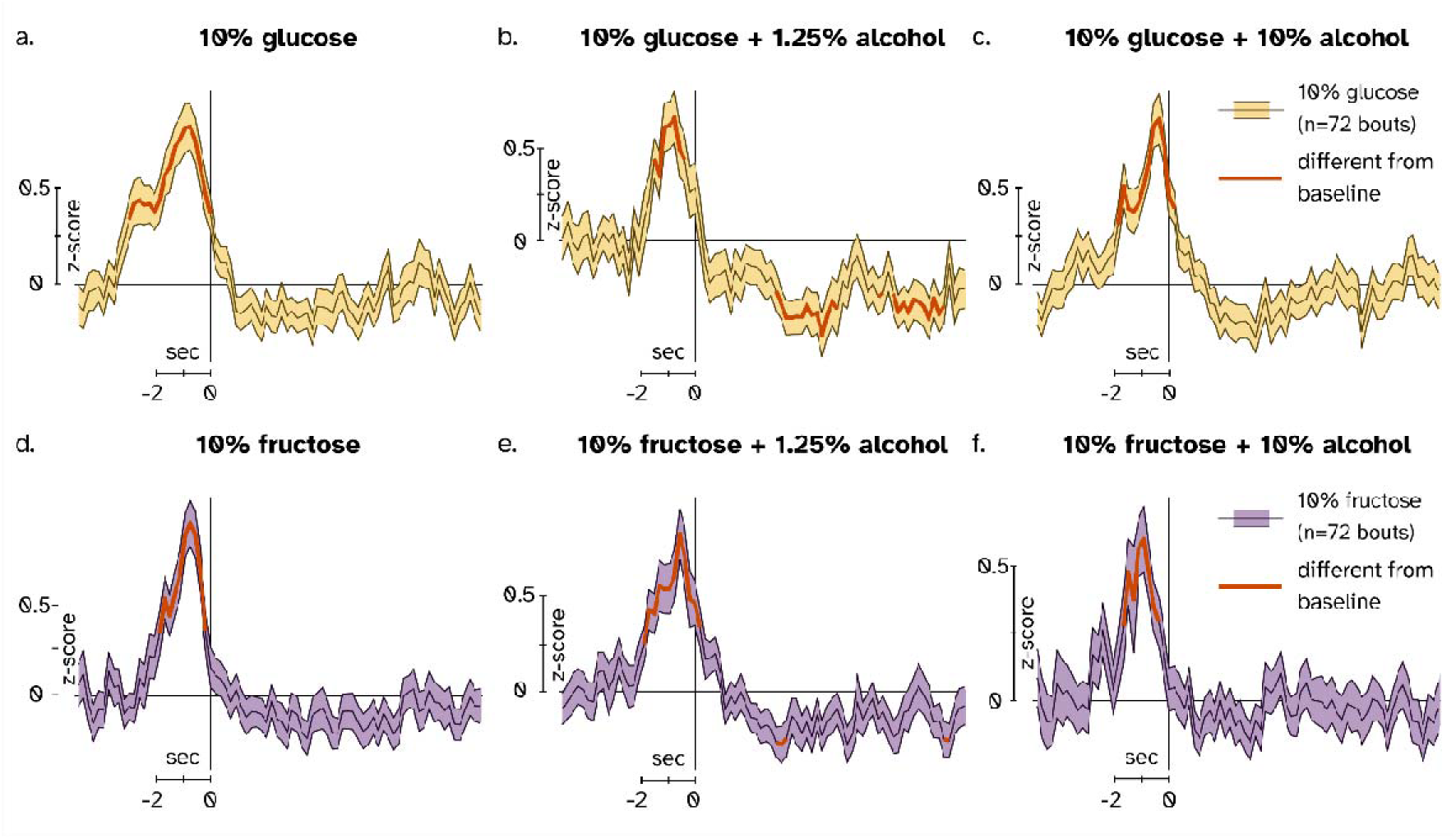
MCH activity peaks before the start of drinking for all cocktails. All data are shown as mean ± SEM. Y-axis scale bar denotes 0.5 z-score of MCH Ca^2+^ activity from baseline, X-axis scale bar denotes 2 seconds. Time intervals where MCH activity is different from baseline (0 z-score) are shown as a darker orange line on the mean (p<0.05). (**a**) MCH activity rose above baseline ∼3 seconds before the start of drinking 10% glucose (x=0), peaked before drinking, and decreased to baseline soon after the start of drinking. (**b**) MCH activity rose above baseline ∼1.5 seconds before the start of 10% glucose + 1.25% alcohol drinking, peaked before drinking, and decreased past baseline until ∼9s after the start of drinking. (**c**) MCH activity rose above baseline ∼2 seconds before the start of 10% glucose + 10% alcohol drinking, peaked before drinking, and decreased lower than baseline until ∼5s after the start of drinking before returning to baseline. (**d**) MCH activity rose above baseline ∼2 seconds before the start of drinking 10% fructose, peaked before drinking, and decreased to baseline soon after the start of drinking. (**e**) MCH activity rose above baseline ∼2 seconds before the start of 10% fructose + 1.25% alcohol drinking, peaked before drinking, and decreased briefly to below baseline at ∼3s and ∼9s after the start of drinking. (**f**) MCH activity rose above baseline ∼1.5 seconds before the start of 10% fructose + 10% alcohol drinking, peaked before drinking, and decreased to baseline before the start of drinking.

See **Supplemental Fig. S2** for a comparison of glucose and fructose solutions with the same alcohol concentration. We then plotted the MCH calcium activity of both the glucose and the fructose cocktails overlaid within the same alcohol concentration (**Supplemental Fig. S1a-c**).

### Cocktail type impacts MCH calcium activity dynamics but not magnitude

We analyzed the peak MCH activity near the start of drinking for each bout across several variables. First, we found that the time of the peak MCH activity varied across alcohol concentration where peak MCH activity was earlier with the higher alcohol concentration (**Fig. 5a**, two-way Mixed Effects Analysis, fixed effect of *alcohol concentration*, *F*_2,227_=4.043, *p*=0.0188), but this change across alcohol concentration depended on the sugar type, where significant changes appeared for only fructose cocktails (*sugar type* x *alcohol concentration* interaction, *F*_2,227_=6.041, *p*=0.0028). Next, we failed to find a significant effect of cocktail type on how long the MCH activity was above baseline (**Fig. 5b**). Similarly, cocktail type was not found to impact the area under the curve of the peak MCH activity and the duration of the peak (**Fig. 5c**). Again, there was no significant effect of cocktail type on the maximum MCH activity (**Fig. 5d**). However, both the sugar and the alcohol concentration of the cocktail affected the MCH activity at the start of drinking, where MCH activity decreased with the high alcohol fructose cocktail (**Fig. 5e**, *sugar type* x *alcohol concentration* interaction, *F*_2,284_=5.015, p=0.0072).

**Fig. 5.**
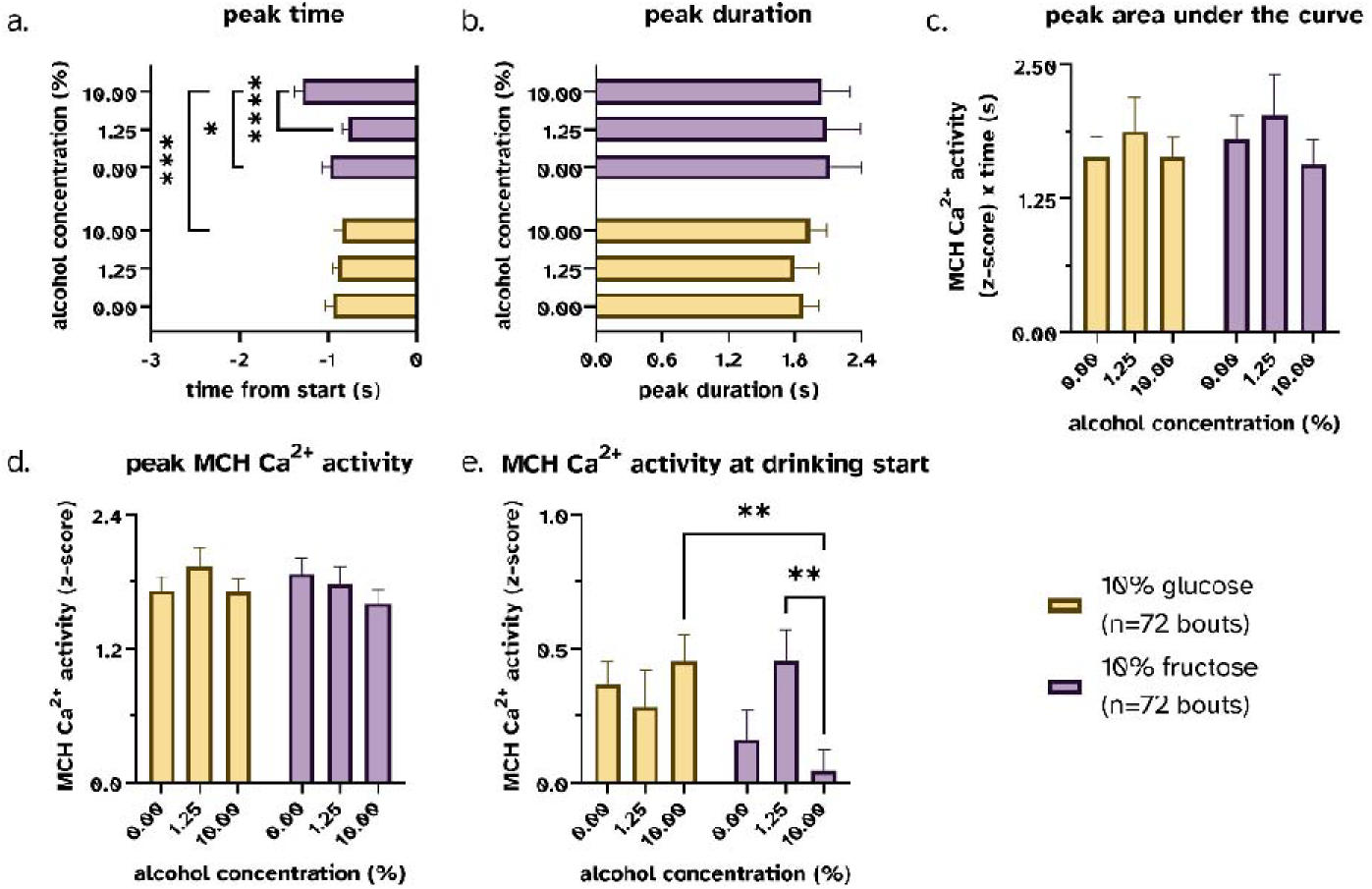
Timing of peak MCH activity varies by sugar type and alcohol percentage. All data are shown as mean ± SEM. (**a**) MCH activity peaked earliest for 10% fructose + 10% alcohol, around 1s before the start of drinking for all glucose cocktails and 10% fructose with no alcohol, and latest for 10% fructose + 1.25% alcohol. (**b**) MCH activity around the start of drinking remained raised for similar durations for each sugar and cocktail solution. (**c**) The area under the curve of raised MCH activity was similar for each sugar and cocktail solution. (**d**) The maximum MCH activity before the start of drinking was similar for each sugar and cocktail solution. (**e**) The MCH activity at the start of drinking was lower for 10% fructose + 10% alcohol compared to 10% glucose + 10% alcohol and compared to 10% fructose + 1.25% alcohol. * p <0.05, ** p<0.01, ***p<0.001, **** p<0.0001

### Peak MCH calcium activity during drinking is correlated with approach behavior

We next analyzed the three timepoints within each cocktail type using the chi-square test of independence to determine whether the timepoints around the start of drinking were correlated with different behaviors. The individual behaviors categorized in each group for the ethological analysis are found in **Supplemental Table S1**. Most behaviors listed were behaviors observed during the course of the experiment. During 10% glucose drinking without alcohol added, each timepoint was correlated with different behaviors (**Fig. 6a**, Χ^2^ (6, *N*=81)=21.64, *p*=0.0014). During 10% glucose + 1.25% alcohol drinking, there were no incidences of drinking-related behaviors such as rhythmic mouth movements; therefore, it was not possible to analyze all behavioral categories together using the chi-square test. The lack of drinking-related behaviors such as rhythmic mouth movements during glucose with low alcohol drinking may be due to its moderate palatability, higher than all the other cocktail types (Kuebler et al. 2026) but likely lower than the sugar solutions.

**Fig. 6.**
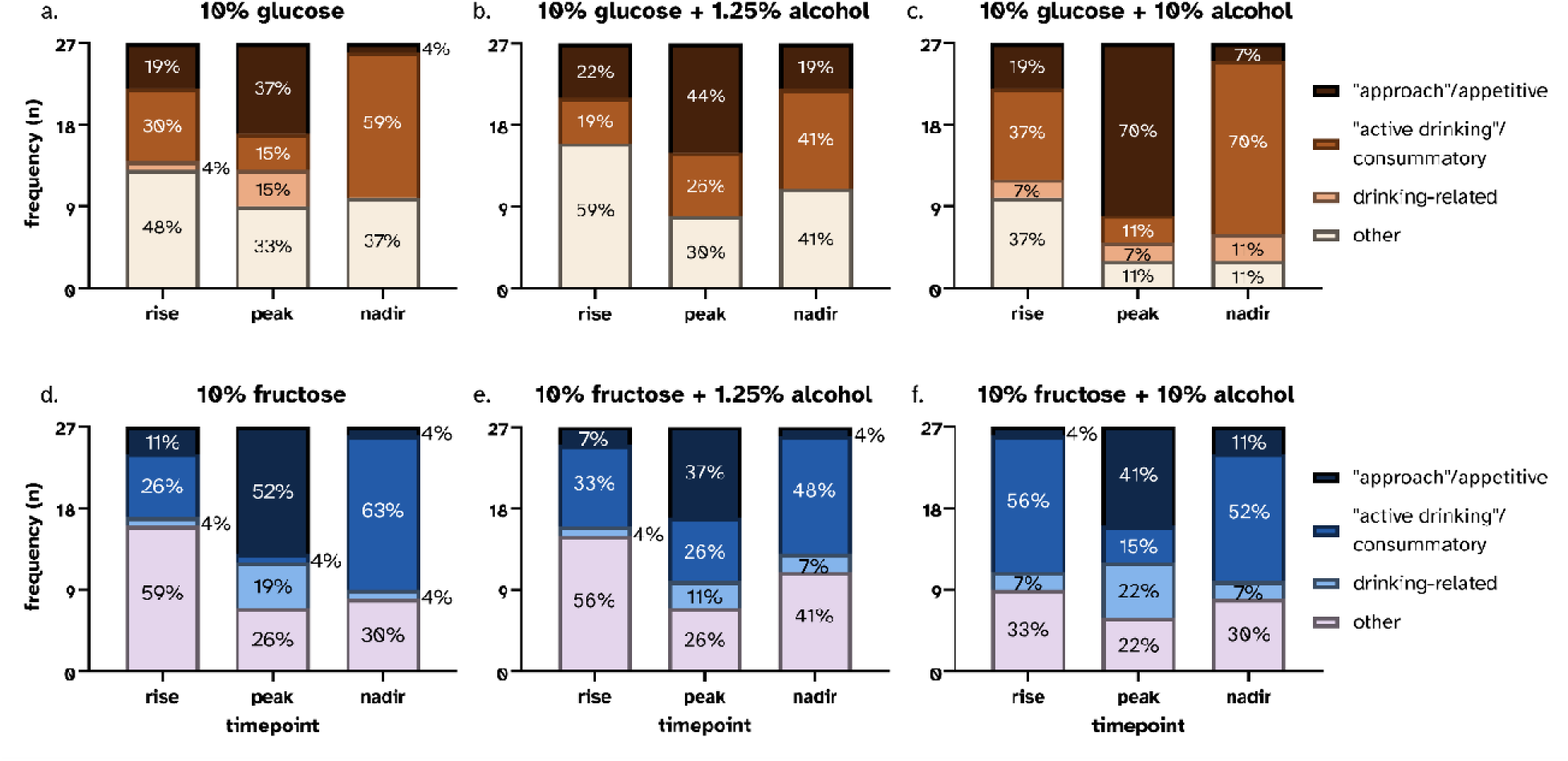
Ethology of female rats drinking sugar and sugar-sweetened cocktail solutions at three timepoints around the start of drinking: the initial rise in MCH activity before the start of drinking, the peak of MCH activity, and the nadir of MCH activity after the start of drinking when MCH activity returns to baseline or dips below baseline. Each set of 3 columns shows the same n=27 drinking bouts. (**a**) While drinking 10% glucose without alcohol, behaviors not related to drinking were most common during the initial rise in MCH activity, appetitive behaviors were most common at the peak of MCH activity, and active drinking was most common when MCH activity rose again after returning to baseline. (**b**) While drinking 10% glucose + 1.25% alcohol, non-drinking-related behaviors were most common during the rise in MCH activity before drinking, appetitive behaviors were most common at the highest MCH activity, and both active drinking and non-drinking-related behaviors were most common during the undershoot of MCH activity. (**c**) With 10% glucose + 10% alcohol drinking, both active drinking and non-drinking-related behaviors were most common during the initial rise in MCH activity, appetitive behaviors were highly prevalent at the peak of MCH activity, and active drinking was highly prevalent during the nadir of MCH activity. (**d**) While drinking 10% fructose without alcohol, non-drinking-related behaviors were most prevalent while MCH activity rises before drinking, appetitive behaviors were most common at the highest MCH activity, and active drinking was the most frequent behavior at the nadir of MCH activity after drinking. (**e**) With 10% fructose + 1.25% alcohol, behaviors not related to drinking were most common at the initial rise in MCH activity, appetitive behaviors were most frequent at the peak of MCH activity, and active drinking was most common during the undershoot of MCH activity. (**f**) With 10% fructose + 10% alcohol, active drinking was the most common behavior while MCH activity started to rise before drinking, appetitive behaviors were most prevalent during the highest MCH activity, and active drinking was most common when MCH activity rises again after returning to baseline.

Chi-square analysis of the three remaining behavioral categories failed to indicate differences. During 10% glucose + 10% alcohol drinking, the three timepoints had independent ratios of behaviors (**Fig. 6b**, Χ^2^ (6, *N*=81)=37.27, *p*<0.0001). For 10% fructose drinking with no alcohol added, each timepoint was correlated with different behaviors (**Fig. 6d**, Χ^2^ (6, *N*=81)=41.29, *p*<0.0001). Similarly, during drinking of 10% fructose + 1.25% alcohol, the behavioral categories were independent among the three timepoints (**Fig. 6e**, Χ^2^ (6, *N*=81)=17.07, *p*=0.009). Lastly, during 10% fructose + 10% alcohol drinking, each timepoint was again correlated with different behaviors (**Fig. 6f**, Χ^2^ (6, *N*=81)=21.74, *p*=0.0014).

Next, we performed exploratory follow-up analyses comparing the incidence of a certain behavioral category during drinking of two cocktail types. For each timepoint, we compared one behavioral category (i.e., appetitive or consummatory) versus all other behaviors between 1.25% and 10% alcohol for each sugar type, and between the two sugar types at 10% alcohol. Results are shown in **Table 1**.

**Table 1.** Fisher’s exact test results of the noted behavior category versus all other behaviors between two cocktail types. * p<0.05, ns = not significant.

| cocktail (more) | cocktail (fewer) | timepoint | behavior category | p-value | significant |
| --- | --- | --- | --- | --- | --- |
| 10% glucose + 10% alcohol<br>Fig. 6c | 10% glucose + 1.25% alcohol<br>Fig. 6b | rise | "active drinking"/<br>consummatory | 0.112 | ns |
| 10% fructose + 10% alcohol<br>Fig. 6f | 10% fructose + 1.25% alcohol<br>Fig. 6e | rise | "active drinking"/<br>consummatory | 0.314 | ns |
| 10% glucose + 10% alcohol<br>Fig. 6c | 10% fructose + 10% alcohol<br>Fig. 6f | rise | "active drinking"/<br>consummatory | 0.950 | ns |
| **10% glucose + 10% alcohol Fig. 6c** | **10% glucose + 1.25% alcohol Fig. 6b** | **peak** | **"approach"/**<br>**appetitive** | **0.049** | **\*** |
| 10% fructose + 10% alcohol<br>Fig. 6f | 10% fructose + 1.25% alcohol<br>Fig. 6e | peak | "approach"/<br>appetitive | 0.500 | ns |
| **10% glucose + 10% alcohol Fig. 6c** | **10% fructose + 10% alcohol Fig. 6f** | **peak** | **"approach"/**<br>**appetitive** | **0.027** | **\*** |
| **10% glucose + 10% alcohol Fig. 6c** | **10% glucose + 1.25% alcohol Fig. 6b** | **nadir** | **"active drinking"/**<br>**consummatory** | **0.027** | **\*** |
| 10% fructose + 10% alcohol<br>Fig. 6f | 10% fructose + 1.25% alcohol<br>Fig. 6e | nadir | "active drinking"/<br>consummatory | 0.500 | ns |
| 10% glucose + 10% alcohol<br>Fig. 6c | 10% fructose + 10% alcohol<br>Fig. 6f | nadir | "active drinking"/<br>consummatory | 0.132 | ns |

During peak MCH activity, there were more incidences of appetitive behaviors while drinking 10% glucose + 10% alcohol (**Fig. 6c**) than 10% glucose + 1.25% alcohol (**Fig. 6b**, Fisher’s exact test, *p*=0.049). This same rate of appetitive behaviors with 10% glucose + 10% alcohol (**Fig. 6c**) was also greater than with 10% fructose + 10% alcohol (**Fig. 6f**, *p*=0.027). During the nadir of MCH activity after the start of drinking, there were more active drinking behaviors during 10% glucose + 10% alcohol drinking (**Fig. 6c**) compared to 10% glucose + 1.25% alcohol drinking (**Fig. 6b**, *p*=0.027).

## Discussion

The objectives of this study were to investigate the interaction between MCH neuronal activity, drinking behavior associated with sugar and sugar-sweetened cocktails, and non-drinking sensory stimuli of different modalities. We sought to determine whether MCH calcium activity was correlated differently with glucose, fructose, and alcoholic cocktail drinking. Further, we determined the timing and magnitude of MCH calcium activity during each stimulus and sugar or cocktail solution. MCH neuronal activity, measured by Ca^2+^ imaging, rapidly rose at the onset of sensory stimuli. In contrast, MCH neuronal activity peaked before the start of drinking for each sugar and cocktail solution. Peak timings of MCH activity varied amongst the two sugar solutions and alcohol concentrations, while peak magnitude remained consistent. Lastly, in female rats, peak MCH activity was correlated with approach behavior for all sugar and cocktail types.

### MCH calcium activity is highly responsive to non-drinking sensory stimuli

Consistent with our hypothesis, MCH calcium activity rose in response to non-drinking, sensory stimuli. However, compared to other measures of *in vivo* LH activity, like glucose electrochemical detection (Kuebler et al., 2022), the MCH calcium activity was unexpectedly pronounced and robust. Specifically, the rise in MCH calcium in response to non-drinking sensory stimuli was rapid and observed consistently across sensory modalities.

Both the novel object and tail touch stimuli (**Fig. 3 a-b**) induced a rise from baseline before the recorded start of the stimulus. That is, when the object contacted the chamber floor, or the moment when the clothespin was secured to the rat’s tail, respectively. This rise is likely due to the rat responding to events preceding the recorded start of the stimuli, which could have included seeing the experimenter approach the chamber through the wall of the transparent chamber, or handling of the rat’s tail just prior to attaching the clothespin to the tail. This initial rise in MCH activity underscores the sensitivity and dynamic responses of these neurons to peri-event stimuli preceding the recorded start of the event. These findings also closely mirror others of MCH activity during a novel object presentation which demonstrated similar dynamics of the initial rise, peak, and return to baseline (Kosse & Burdakov, 2019).

Extracellular glucose in the lateral hypothalamus also rises at the onset of a novel object presentation (Kuebler et al., 2022), which may result from hyperemic increases coupled to local neural activity (Fellows & Boutelle, 1993; Hoge et al., 2005; Kiyatkin & Lenoir, 2012). In our prior work, extracellular glucose levels rose rapidly after the start of the novel object stimulus and then rapidly returned to baseline, similar to the timescale of MCH calcium activity (Kuebler et al., 2022). For example, we found that glucose levels rose at the start of the tail touch stimulus and returned to baseline levels about 20s later. The current experiment extend those previous results in two ways. First, the current findings indicate that changes in lateral hypothalamic extracellular glucose levels in response to naturalistic stimuli are consistent with changes in neuronal activity in the lateral hypothalamus. Second, the current findings suggest that this purported neuronal activity is likely due to increased MCH neuronal activity. However, future research leveraging more causal methods are necessary to directly assess coupling between the activity of MCH neurons directly assessed here and lateral hypothalamic glucose levels.

Moreover, there is a temporal difference between MCH calcium activity and lateral hypothalamic extracellular glucose levels, with a delay of 3-4s between sensory stimuli and cerebral blood flow changes (Enager et al., 2009; Zimmer et al., 2011). Determining whether these differences in timing result only from differences in monitoring methodology or also from differences in physiological processes, such as contributions from other neuronal systems in the lateral hypothalamus, will need to be investigated further. Together, the high magnitude MCH responses to a wide variety of sensory stimuli suggests that MCH neurons are the primary neuronal system integrating fast sensory inputs in the lateral hypothalamus.

### MCH calcium activity peaks before the start of drinking for all sugar and cocktail solutions

As expected, MCH activity quickly rose during free drinking behavior. Surprisingly, MCH calcium activity peaked before the start of drinking and returned to baseline close to the start of drinking. Further, rats drinking both glucose and fructose cocktails containing a low alcohol concentration also showed MCH calcium activity below baseline levels several seconds later, often when they were still drinking.

Our findings contribute to growing evidence for MCH activity during both the appetitive phase of ingestive behavior before active feeding or drinking as discussed here, and during the consummatory phase of active feeding or drinking (for review see Kongstorp et al., 2025). In terms of appetitive food-seeking behaviors, MCH activity is increased in response to food-predictive cues and during food-seeking behaviors (Subramanian et al., 2023). In a subset of female rats in our study, we found increased MCH activity during approach before drinking.

These results suggest that MCH activity may regulate appetitive behaviors for sugar and sweetened cocktail drinking, though the precise mechanism of this regulation requires future research.

In terms of consummatory behaviors, MCH responses may differ depending on the mode of ingestion (i.e., eating versus drinking). For example, similar to our findings, others have reported decreased MCH activity after the onset of drinking (Concetti et al., 2024). In contrast, during chow feeding MCH calcium activity has been shown to increase compared to non-feeding periods, particularly during early feeding bouts (Subramanian et al., 2023). Moreover, optogenetic stimulation of MCH neurons during active feeding increases consumption (Dilsiz et al., 2020). Thus, it appears that there are consistent and important differences in the dynamics and direction of MCH neuronal responses to eating and drinking consummatory behaviors.

Future work will need to parse whether the dynamic effects of MCH calcium activity in our study, that is, the quick rise and fall before the start of drinking, are unique to alcoholic cocktails in rats or are more generalizable to other types of drinking. Interestingly, the phasic dynamics of MCH calcium activity during drinking have some similarities with the firing characteristics of some GABAergic medium spiny neurons in the nucleus accumbens (Nicola et al., 2004), a mesocorticolimbic structure that also receives abundant input from lateral hypothalamic MCH neurons. This similarity suggests that MCH projections to the nucleus accumbens may contribute to mesolimbic responses to reward predictive cues.

It should also be noted that the peak height of MCH activity before the start of drinking did not appear to reach a ceiling, further supported by the observation that MCH activity during non-drinking stimuli compared to drinking was much higher (**Fig. 3 a, b, d**). Importantly, we did not normalize the MCH calcium signal across subjects or sessions statistically. Therefore, the peak heights across cocktail type were consistent as observed, and the height of peak MCH activity was not obscured by normalization (Wallace et al., 2025).

### MCH activity dynamics vary across cocktail type

Because we observed that the MCH signal peaked before the start of drinking, we sought to focus on investigating timing differences in MCH activity among the sugar and cocktail solutions. We were also able to evaluate differences in the magnitude of MCH activity across each cocktail type by directly comparing MCH calcium signal peak height and its related measure area under the curve. Surprisingly, we did not find effects related to the magnitude of peak MCH activity across cocktail type. Therefore, any effects of cocktail type and MCH activity are not likely resultant from recruiting more MCH neurons during peak activity. Rather, greater excitability of MCH neurons in response to cocktail contents may raise baseline MCH levels. Yet, this speculation requires confirmation using methods suited to recording slower baseline changes in neuronal activity.

Regarding timing effects, our findings show that alcohol concentration impacted the dynamics of MCH calcium activity for fructose cocktails, but not glucose cocktails (**Fig. 5a, e**). That is, increasing alcohol in the brain affected MCH activity when drank in cocktails with minimal sugar entry into the brain, such as with fructose. Still, the mechanisms for how different cocktail types may impact MCH dynamics are unclear.

First, one way that MCH neurons may discriminate between cocktail types is by glucose sensing, that is, increasing in excitability in response to greater extracellular glucose concentrations (Burdakov et al., 2005). Glucose concentrations quickly rise in the brain and peak around 15-20 minutes following drinking glucose solutions (Wakabayashi & Kiyatkin, 2015). In the current study, drinking sessions were ∼25 minutes long, meaning that drinking bouts in the first ∼20 minutes of the session occurred while glucose levels are rising in the brain. To control for the effect of glucose or alcohol entering the brain, we used the same number of drinking bouts from the beginning, middle, and last bouts of the drinking session. Most middle and late drinking bouts occurred several minutes or more (∼4 and ∼13 min) after the first drinking bouts. Therein, MCH activity should be interpreted as reflecting, on average, the still rising glucose and alcohol levels in the brain. As there were large differences among the intervals between early and late bouts, as well as amounts of cocktails drank, we did not compare early and late bouts. Investigating MCH activity after certain intervals of drinking, such as longer drinking sessions to compare rising glucose levels with high glucose levels after ∼30 minutes, will be valuable for understanding how MCH responds to glucose in the brain.

Particularly, it will be informative to examine hypothesized effects of glucose that occur very early in the drinking session (Davis, 1989; Schier & Spector, 2016) alongside behavioral measures of post-ingestive effects (Kuebler et al. 2026).

Second, alcohol, like glucose, is well-known for affecting various brain functions. Interestingly, our findings indicated that increasing alcohol concentration impacted MCH dynamics for fructose cocktails but not for glucose cocktails (**Fig. 5a, e**). One hypothesis for why alcohol may affect MCH calcium activity for fructose but not glucose drinking is by an interaction of the central effects of the cocktail and its palatability. It may be that rising alcohol concentrations promote alcohol drinking despite the aversive taste of alcohol by directly activating MCH neurons. With fructose cocktails, low alcohol concentrations may promote earlier drinking after the MCH peak, while higher alcohol concentrations may delay drinking, which still results in greater alcohol intake than lower alcohol cocktails. However, glucose in cocktails may mask the dose-dependent effects of alcohol by activating MCH neurons regardless of alcohol concentration. To wit, we observed that both glucose cocktails resulted in drinking soon after the MCH peak.

Lastly, how alcohol affects MCH neuron activity remains unclear and understudied. On the one hand, alcohol may inhibit MCH neurons via conventional receptor binding, such as through GABA_A_ or multiple glutamatergic receptors (Monti et al., 2013; Yang et al., 2022).

However, MCH signaling promotes alcohol drinking (Cippitelli et al., 2010; Karlsson, Rehman, et al., 2016), and acute alcohol exposure stimulates neuronal MCH expression (Morganstern, Chang, Chen, et al., 2010), suggesting that acute alcohol likely activates MCH neurons to promote further alcohol drinking. Local network input from orexin neurons may also promote MCH expression and activity (Morganstern, Chang, Chen, et al., 2010), though the mechanism for stimulation of orexin is unclear (Morganstern, Chang, Barson, et al., 2010). Another hypothesis is that alcohol may directly increase MCH activity. This process could be via an unknown excitatory receptor, or as a type of nutrient sensing via alcohol itself or of acetate, a metabolite of alcohol, similar to glucose sensing (Kuebler et al. 2026). This potential unknown nutrient sensing mechanism also indicates further interactions between alcohol and glucose.

For example, alcohol decreases glucose metabolism and promotes acetate metabolism in many areas of the brain (Volkow et al., 2013). Further, the hypothalamus metabolizes acetate from the diet more than other brain areas, affecting the activity of other neuropeptide systems in the lateral hypothalamus (Forte et al., 2024) via indirect metabolism by glia (Frost et al., 2014).

Taken together, regardless of its exact mechanism of action, our findings suggest that alcohol has more of an activating effect on MCH neurons than previously known. Our study is the first to investigate MCH activity during alcoholic cocktail drinking, and more investigation of MCH activity during both sugar and alcohol drinking will provide essential information for understanding how MCH neurons regulate drinking.

One hypothesis of how MCH regulates feeding is that MCH neurons facilitate associative learning between the taste of nutritive food and raised brain glucose levels. This direct motivational component of MCH activity is thought to be via MCH release in mesolimbic areas from the lateral hypothalamus concurrently with mesolimbic input from taste centers (Domingos et al., 2013). Others have suggested that the same taste stimulus then acts as a conditioned stimulus after associative learning to promote feeding preferences for glucose-containing food (Kosse et al., 2015). Namely, the activation of MCH neurons shifts to conditioned stimuli associated with the unconditioned reward. We found a dynamic rise in MCH activity during approach behavior but not during drinking, which supports a role for MCH activity in conditioned responses to cocktail-related cues. In our study, rats were trained to drink before the start of recording sessions, so all rats were poised to freely associate drinking-related cues. In following recording sessions, there were many sensory cues linked to drinking present before the start of drinking. Both the visual cue of the nozzle, which was the same between glucose and fructose solutions, and the smell of the cocktail, which likely varied depending on the sugar, were present during approach behaviors. Here, if MCH neurons are active concurrently with glucose-associated sensory cues as assumed, then our findings indicate that MCH may respond to more than taste cues alone. MCH neurons also integrate olfactory inputs to help regulate feeding (Adams et al., 2011), so it is plausible to suggest that visual or olfactory cues could provide the same kind of conditioned stimulus as those from taste (Domingos et al., 2013). How MCH neurons integrate conditioned stimuli from multiple modalities will need to be determined empirically to understand its role in ingestive behaviors with glucose sensing.

### Maximum MCH calcium activity is correlated with appetitive behavior for each sugar and cocktail solution in female rats

We also examined the proportions of appetitive, consummatory, drinking-related, and other behavior during the peak of MCH activity. These were also investigated during the initial rise before drinking and the nadir after drinking when MCH activity began to rise again. During sugar and cocktail drinking, peak MCH calcium activity was most strongly associated with appetitive behaviors such as approaching or reorienting toward the nozzle for female rats. This is in contrast to prior findings of MCH calcium activity during chow feeding, from which activity was associated with both appetitive and consummatory behaviors in male rats (Subramanian et al., 2023). Future work will need to determine which variables impact the associations between MCH activity and ingestive behaviors, including sex differences and type of behavior, such as drinking or feeding. For both sugar solutions and both cocktails containing a low alcohol concentration, the initial rise in MCH activity was correlated with behaviors not directly related to drinking (**Fig. 6a-b, d-e**). In other words, MCH activity rose before the rats approached the nozzle. This study found behaviors associated with the peak of MCH activity. The reverse analysis, finding the MCH activity associated with transitions between behaviors, may also be fruitful (Mahler & Berridge, 2009). However, for cocktails containing a high alcohol concentration, the rise in MCH activity was equally or more correlated with active drinking (**Fig. 6c, f**).

As expected, active drinking was the most common behavior at the nadir, immediately before MCH activity rose again after the start of drinking. Our definition of drinking bouts included a 1s pause between bouts. This was shorter than the interval between the start of drinking and the nadir, so it is likely that the rise in MCH activity after the start of drinking may reflect the initial rise before the next drinking bout.

The greatest magnitude difference in rates of behaviors during the MCH peak was found between high alcohol glucose and fructose cocktails. Rats drinking glucose cocktails were very likely to approach the nozzle during MCH peak activity, while rats drinking fructose cocktails were more likely to have remained at the nozzle from a previous drinking bout. Thus, MCH activity was related to approach behaviors more often with glucose in the brain than without glucose. This pattern suggests that glucose in the brain may specifically facilitate approach behaviors, mediated by MCH neurons.

### Limitations and future directions

Our experiments were correlational and suggested that phasic MCH activity may be a mechanism for regulating appetitive behaviors during sugar and cocktail drinking. However, this mechanism requires experimental confirmation, such as determining the impacts of MCH stimulation or inhibition during drinking behavior. This study did not investigate different subpopulations of MCH neurons in the lateral hypothalamus that co-release other neurotransmitters (Mickelsen et al., 2017) or projections to specific brain areas (Chee et al., 2015), which will be necessary to understand the MCH system’s regulation of appetitive and consummatory behaviors (Kongstorp et al., 2025). This experiment focused on the start of drinking and sensory stimuli. However, it may be fruitful to investigate MCH activity at the end of the same stimuli and at the end of drinking using more precise methodology such as lickometers.

This study used both female and male subjects to develop experimental procedures, as MCH fiber photometry data has only been published using female subjects once before (Wang et al., 2024). Future studies on MCH will need to systematically investigate sex as a biological variable (Kuebler et al., 2024). Further highlighting the need for systematic study, the ethology portion of this study was conducted using female rats only. We predict that male rats would have similar results as female rats, but it is possible that male rats would respond differently to glucose and alcohol entering the brain (Kuebler et al., 2024; Mogi et al., 2005).

### Translational implications

This study provides evidence that MCH neurons may regulate cocktail drinking depending on the type of sugar in the cocktail. Our findings suggest that glucose and alcohol concentration interact to stabilize MCH neuronal activity, likely leading to altered regulation of cocktail drinking compared to fructose cocktails. Therefore, this study provides evidence that glucose and alcohol within sweetened cocktails impact neuronal activity in the brain, possibly altering drinking outcomes. Additionally, our study found that heightened MCH activity is correlated with the appetitive phase of sugar and cocktail drinking. As this phase of drinking is critical in sugar and cocktail seeking, MCH may be an important target for limiting alcohol drinking and uncovering the biological mechanisms that contribute to AUD.

## Supplemental material

Supplemental data is available at *Figshare* data:10.6084/m9.figshare.32003715

## Data Availability

All data are available in the main text and supplemental data. Raw data supporting the conclusions of this article will be made available upon request.

## Acknowledgments

The authors would like to acknowledge the technical assistance of Dr. You (Joe) Zhou (Morrison Microscopy Core), who has provided confocal microscopy assistance for this manuscript.

## Author contributions

**I.R.K.K**: Conceived and designed research; Analyzed data; Performed experiments; Interpreted results of experiments; Prepared figures; Drafted manuscript; Edited and revised manuscript; Approved final version. **J.C**. Analyzed data; Performed experiments: **J.V**. Analyzed data; Performed experiments, **C.E.B**: Conceived and designed research; Performed experiments; Edited and revised manuscript, **N.A.H**.: Edited and revised manuscript, **K.T.W**.: Conceived and designed research; Performed experiments; Interpreted results of experiments; Edited and revised manuscript; Approved final version.

